# Parkour LIMS: facilitating high-quality sample preparation in next generation sequencing

**DOI:** 10.1101/338533

**Authors:** E. Anatskiy, D.P. Ryan, B. Grüning, L. Arrigoni, T. Manke, U. Bönisch

## Abstract

**Summary:** This paper presents Parkour, a software package for sample processing and quality management of next generation sequencing data and samples. Starting with user requests, Parkour allows tracking and assessing samples based on predefined quality criteria through different stages of the sample preparation workflow. Ideally suited for academic core laboratories, the software aims to maximize efficiency and reduce turnaround time by intelligent sample grouping and a clear assignment of staff to work units. Tools for automated invoicing, interactive statistics on facility usage and simple report generation minimize administrative tasks. Provided as a web application, Parkour is a convenient tool for both deep sequencing service users and laboratory personal. A set of web APIs allow coordinated information sharing with local and remote bioinformaticians. The flexible structure allows workflow customization and simple addition of new features as well as the expansion to other domains.

**Availability and implementation:** The code and documentation are available at https://github.com/maxplanck-ie/parkour

## Introduction

Next generation sequencing (NGS) technology and applications are rapidly evolving, leading to an ever increasing number of samples requiring processing. Upstream of sequencing and data analysis, sample batches undergo a series of interdependent preparatory steps that need to be conducted in a precise and timely manner. Moreover, these processes also generate large amounts of metadata, which can be important for follow-up analyses and comparative studies. Typically, core facilities support technologically advanced and data intensive projects and rely on Laboratory Information Management Systems (LIMS) for documentation and tracking. Specifically for NGS applications many commercial or open source LIMS exist that focus on different aspects of the workflow. BaseSpace Clarity LIMS (GenoLogics) is a complete commercial, highly customizable solution, though with a high license fees hindering a widespread use, especially in mid-sized academic laboratories. Multiple non-commercial open-source solutions exist that offer a wide range of functionality and are applied in laboratories of different scopes and scale. For instance, GNomEx <u>(Nix et al. 2010)</u>, openBIS <u>(Barillari et al. 2016)</u> and SMITH <u>(Venco et al. 2014)</u> provide extensive solutions for sample submission, tracking, billing, data organization and analysis. The data analysis platform Galaxy has a LIMS extension <u>(Scholtalbers et al. 2013)</u> for sample tracking, interactive flowcell design and automated sample demultiplexing. The Wasp System <u>(McLellan et al. 2012)</u> and SMITH <u>(Van Rossum et al. 2010)</u> represent a LIMS preconfigured to support special applications like the analysis of clinical samples or sample management in genotyping laboratories.

However, most of the existing LIMS are missing laboratory notebook aspects to support, organize and ultimately standardize initial laboratory-intensive sample preparation steps.

To this end, we developed Parkour LIMS, which is ideally suited to facilitate and maintain high quality and efficient sample preparation. Both user and laboratory staff have controlled access to the system for maximal transparency and communication. Laboratories, dealing with NGS samples processing, can benefit from using Parkour LIMS as electronic laboratory book and to assist in team planning for standardized NGS sample preparation. Parkour LIMS also facilitates automated sample processing and interaction with bioinformaticians through a variety of web APIs.

## Results

### System Overview

The goal was to implement a database that supports in depth the laboratory work for NGS sample preparation to minimize error rates and maximize reproducibility, cost effectiveness and minimize turnaround time. Furthermore, modifications to account for changing protocols and technologies are possible for any Parkour administrator without specific programming skills. Upon sample submission and approval Parkour guides the request through multiple stages of the NGS workflow, including incoming quality control, index generation, sample preparation, sample pooling and loading of samples onto sequencing instruments (Figure 1).

**Figure 1.**
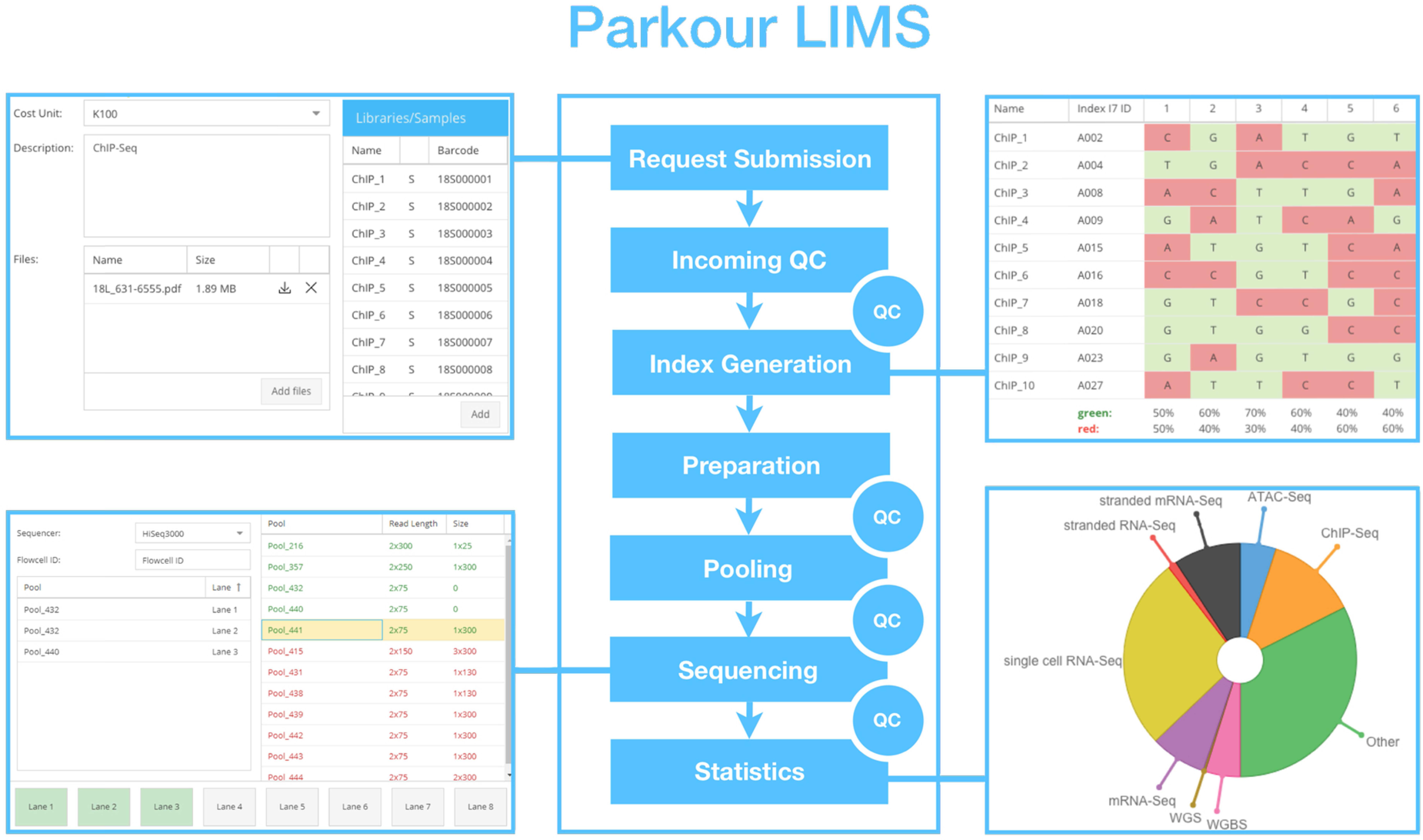
Parkour LIMS workflow. Middle: NGS workflow (QC: quality controls). Top left: Request submission window in Parkour. Top right: Index generation with Parkour. Bottom left: Interactive flowcell loading. Bottom riight: Sample statistics.

A request holder can at any time follow request status and view results of conducted measurements. Parkour coordinates the work of the laboratory staff by provision of clearly structured to-do lists. The software was designed to support laboratory work and documentation as conveniently as possible and to overcome any need for a laboratory notebook. In each stage of the workflow associated samples are displayed and the laboratory staff can edit predefined fields. For batch processing, multiple measurement values can be easily pasted into the forms. In multiple stages, spreadsheets, filled with per-stage relevant information and calculations, can be downloaded and function as templates for further steps (e.g. to instruct adjustment of loading concentrations for sample preparation, pooling or instrument loading). Simple adjustments to calculations can be made, if needed. Raw data from measurements can easily be uploaded to each request for documentation and to share with the request owner. To complete a stage and move samples to the next step, the laboratory staff actively evaluates the results and approves or marks samples as failed for the next step. Parkour has an email notification feature to contact a request holder.

Flexible configuration provided by Parkour enables authorized laboratory staff, without programming skills, to independently manage Parkour user registration and workflow configuration. Sample preparation protocols, predefined list of available indices, lane or run capacities, instrumentation for quality control and sequencing as well as price lists, cost centers and laboratories of principal investigators can be quickly adjusted to minimize interruptions and system downtimes while waiting for code changes by IT staff.

The developed system consists of two principal layers - request submission, carried out by the user, and request processing, executed by the laboratory staff.

### Request submission

Request submission is initiated by filling in a request creation form, where the user adds all the necessary and standardized information about the samples, such as read length, sequencing depth, preparation protocol as well as purity measurements. To ensure data integrity, user input is validated against a controlled vocabulary. Invalid fields will be marked and a message then provides information on expected values for a particular field. Users are also able to upload additional data, quality plots or any other relevant files that are necessary for proper sample processing. After creating a request, all submitted samples receive automatically generated barcodes. Upon request submission and approval, the laboratory staff is notified and can proceed with request processing.

### Request processing

#### Incoming Quality Control

Each request must pass predefined quality criteria. Upon request submission Parkour displays samples or libraries in the incoming quality control window, to be evaluated by staff of the laboratory. Once quality measurements are documented and data is analyzed, samples/libraries either receive approval or are rejected.

#### Index Generator

High-throughput sequencing typically involves multiplexing, which allows the sequencing of multiple samples simultaneously in a single run. Because this step is of central importance, especially in a facility setting with thousands of different samples, Parkour has a customizable index generator to simplify this logistic hurdle. This tool groups samples by compatible index types and run conditions and assigns generated indices to them. The grouped samples are saved as a pool with a running number. The index generator chooses indices from predefined lists to ensure proper image registration on Illumina sequencing devices. Furthermore, the index generator can handle different index types and formats such as single‐ and dual-indexing in individual tubes or standard 96-well plates. For high throughput preparation a start position and direction can be chosen and the index generator will suggest subsequent indices to be assigned to the samples.

#### Sample Preparation and Pooling

Sample preparation and pooling are labor intensive and time consuming steps that benefit from clear organization. A “to-do-list” and preconfigured spreadsheets that function as benchtop protocols guide the laboratory staff through sample preparation and pooling. Samples that rely on identical preparation protocols are merged to ensure optimal utilization of installed preparation devices (i.e., high throughput liquid handling systems). A red/green color code indicates whether pools are ready to be prepared and qualified prior to instrument loading.

#### Loading of Samples

Qualified pools are loaded on flowcells of appropriate sequencing instruments using an interactive flowcell layout. Parkour automatically generates standardized sample sheets that can be used by an external demultiplexing pipeline after sequencing.

#### Run Monitoring and Reporting

Upon completion of sequencing, a number of parameters key to to judging if a run and sequences are within specification are displayed. A reporting function enables compilation of a PDF report that summarizes details of the quality control, preparation and sequencing. The completed report is attached to each request and shared with the user.

#### Administrative tasks

Parkour supports automatic billing of individual requests. Billing is based on predefined, editable prices for quality control, sample preparation and sequencing. Reports on usage of the facilities workflows and instruments are provided as interactive graphs and can be simply downloaded for standardized presentation and reporting.

## Technical Details

Parkour is a web-based extensible LIMS that uses the latest technologies to easily maintain and develop the system further. We integrated Parkour LIMS with our current scripts and workflows to provide an even higher level of automation. Parkour LIMS has three tiers: the client side (front-end), the application (back-end) server, and the database server. These components can be deployed and set up separately or together using Docker and docker-compose. The client side is completely written in JavaScript utilizing the Ext JS framework, which provides a powerful table component (with a fast search, per column filtering, grouping, summary, etc.). Since the system operates as a web application, it can be used simultaneously by multiple users on any computer with a web browser. The installation steps and the API are described in detail in the extensive documentation.

## Conclusions

Parkour focuses on high quality and efficient sample preparation in a next generation sequencing environment. An adaptive, modular structure guides the laboratory staff through the complete sample-to-sequence workflow. Features like the index generator ensure optimal usage of sequencing capacity for timely and cost-efficient sample processing. A billing module liberates lab personal from tedious organizational tasks. The system is designed to allow simple adjustments. To this end, any user with sufficient permission can simply configure the workflow as needed, such as by addition of new library preparation protocols, price lists or index types. Academic laboratories that employ next generation sequencing will profit from this lightweight software solution to organize sample preparation and sequencing. Beyond next generation sequencing applications, Parkour can easily be extended with new features to support different techniques and workflows.

## Acknowledgements

We would like to thank Diana Santacruz and Nadia Kress for critical assessment of the laboratory work considering features of the Parkour LIMS software.

